# Assembly of bacterial cell division protein FtsZ into dynamic biomolecular condensates

**DOI:** 10.1101/2020.08.27.271288

**Authors:** Miguel Ángel Robles-Ramos, Silvia Zorrilla, Carlos Alfonso, William Margolin, Germán Rivas, Begoña Monterroso

## Abstract

Biomolecular condensation through phase separation may be a novel mechanism to regulate bacterial processes, including cell division. Previous work revealed FtsZ, a protein essential for cytokinesis in most bacteria, and the *E. coli* division site selection factor SlmA form FtsZ∙SlmA biomolecular condensates. The absence of condensates composed solely of FtsZ under the conditions used in that study suggested this mechanism was restricted to nucleoid occlusion or SlmA-containing bacteria. Here we report that FtsZ alone can demix into condensates in bulk and when encapsulated in synthetic cell-like systems. Condensate assembly depends on FtsZ being in the GDP-bound state and on crowding conditions that promote its oligomerization. FtsZ condensates are dynamic and gradually convert into FtsZ filaments upon GTP addition. Notably, FtsZ lacking its C-terminal disordered region, a structural element likely to favor biomolecular condensation, also forms condensates, albeit less efficiently. The inherent tendency of FtsZ to form condensates susceptible to modulation by physiological factors, including binding partners, suggests that such mechanisms may play a more general role in bacterial cell division than initially envisioned.

## Introduction

Biomolecular condensation is emerging as an important regulatory mechanism involved in the function and organization of proteins in a wide variety of systems, but its role in bacterial cells has not been appreciated until relatively recently (Azaldegui *et al*, 2020). The dynamic compartments formed through this mechanism, which lack lipid membranes and are in contact with the surroundings from which they physically separate, accumulate specific molecules while excluding others. From an increasing number of studies of mainly eukaryotic proteins, it seems that condensation is favored by macromolecular crowding, protein multivalency and unstructured domains (Banani *et al*, 2017; Shin & Brangwynne, 2017). Nucleic acids are often present in these assemblies and they generally enhance the tendency of proteins to form them, probably because of the associated multivalency of the complexes (Ambadipudi *et al*, 2017; Lin *et al*, 2015; Molliex *et al*, 2015).

There have been relatively few studies dealing with condensates involving bacterial proteins (Abbondanzieri & Meyer, 2019; Al-Husini *et al*, 2018; Heinkel *et al*, 2019; Ladouceur *et al*, 2020; Wang *et al*, 2019). Among them, we recently described condensates (Monterroso *et al*, 2019) involving FtsZ, a key protein whose polymers organize into a dynamic ring-like structure required for bacterial division (Haeusser & Margolin, 2016), and SlmA, a DNA-binding protein that blocks FtsZ rings assembly over nucleoids in *E. coli* through direct interaction with FtsZ (Mannik & Bailey, 2015; Schumacher, 2017). On the basis of this finding, we proposed that biomolecular condensation may play a role in the regulation of bacterial division, through the modulation of nucleoid occlusion by SlmA (Monterroso *et al.*, 2019).

FtsZ, one of the most conserved proteins across bacterial species, is the central protein of the cytokinesis machinery. From a structural point of view, FtsZ monomers contain a few disordered amino acids at their extreme N-terminus, followed by a globular domain, an unstructured flexible linker (~50 amino acids) and a conserved C-terminal tail (Erickson *et al*, 2010). The GDP bound protein forms non-cooperative isodesmic oligomers, through a process enhanced by low salt and high magnesium concentration (Rivas *et al*, 2000). These oligomers become larger in crowding conditions (Rivas *et al*, 2001) and it is under these circumstances when, upon binding to SlmA, FtsZ organizes into dynamic biomolecular condensates, enhanced by DNA carrying the specific SlmA binding sites (SBS, (Monterroso *et al.*, 2019)). In the presence of GTP and following a cooperative mechanism (Du & Lutkenhaus, 2019), FtsZ associates into filaments of different sizes and morphologies that remain assembled until GDP accumulates as a consequence of GTP hydrolysis.

No biomolecular condensates or any other kind of structure detectable by confocal fluorescence microscopy or turbidity have been found with FtsZ on its own, under the experimental conditions assayed in the aforementioned study. To the best of our knowledge, biomolecular condensates containing FtsZ have not been reported elsewhere. However, as condensation has been frequently associated with multivalency and unstructured regions, it seems that FtsZ itself would be a good candidate for this type of behavior. Furthermore, the plasticity of FtsZ, reflected in its ability to self-associate into different structures depending on experimental parameters, suggests that FtsZ condensates might form under a particular set of conditions not previously tested. This behavior may have been unnoticed in other crowding studies with FtsZ, as most of them have focused on the effects on the GTP-triggered polymers.

Here we have analyzed whether FtsZ itself could undergo biomolecular condensation. We have evaluated the effects of protein and magnesium concentration, ionic strength, and crowding on the ability of FtsZ to form structures detectable by turbidity and fluorescence microscopy, either in bulk or when encapsulated in cytomimetic containers generated by microfluidics. We found structures ranging from irregular patches to round condensate-like arrangements, often coexisting, under conditions promoting oligomerization of the protein in its GDP-bound form. FtsZ condensates were observed only under a narrow range of conditions; removal of the unstructured linker of the protein did not preclude their formation but reduced the extent of their condensation. These results indicate that the main constituent of the bacterial division ring, FtsZ, has the intrinsic ability to form biomolecular condensates.

## Results

### GDP bound FtsZ forms micrometer-sized condensates in crowding conditions

Following a stepwise strategy (**Supplementary Fig. S1**), we tested the ability of FtsZ to form biomolecular condensates on its own by exploring conditions known to promote its oligomerization, such as high magnesium and crowder concentrations, and low salt concentration. Confocal images of FtsZ labeled with Alexa 488 (FtsZ-Alexa 488) showed that, at 100 mM KCl, 10 mM Mg^2+^ and in the presence of dextran, Ficoll or PEG as crowders, the protein (5 µM) formed round structures that resembled biomolecular condensates (**Fig. 1a**). Turbidity measurements on these samples further confirmed the formation of higher order structures in the presence of crowding agents (**Fig. 1b**), with absorbance values in the absence of crowders virtually zero (0.002 ± 0.002 with 16 μM FtsZ).

**Figure 1.**
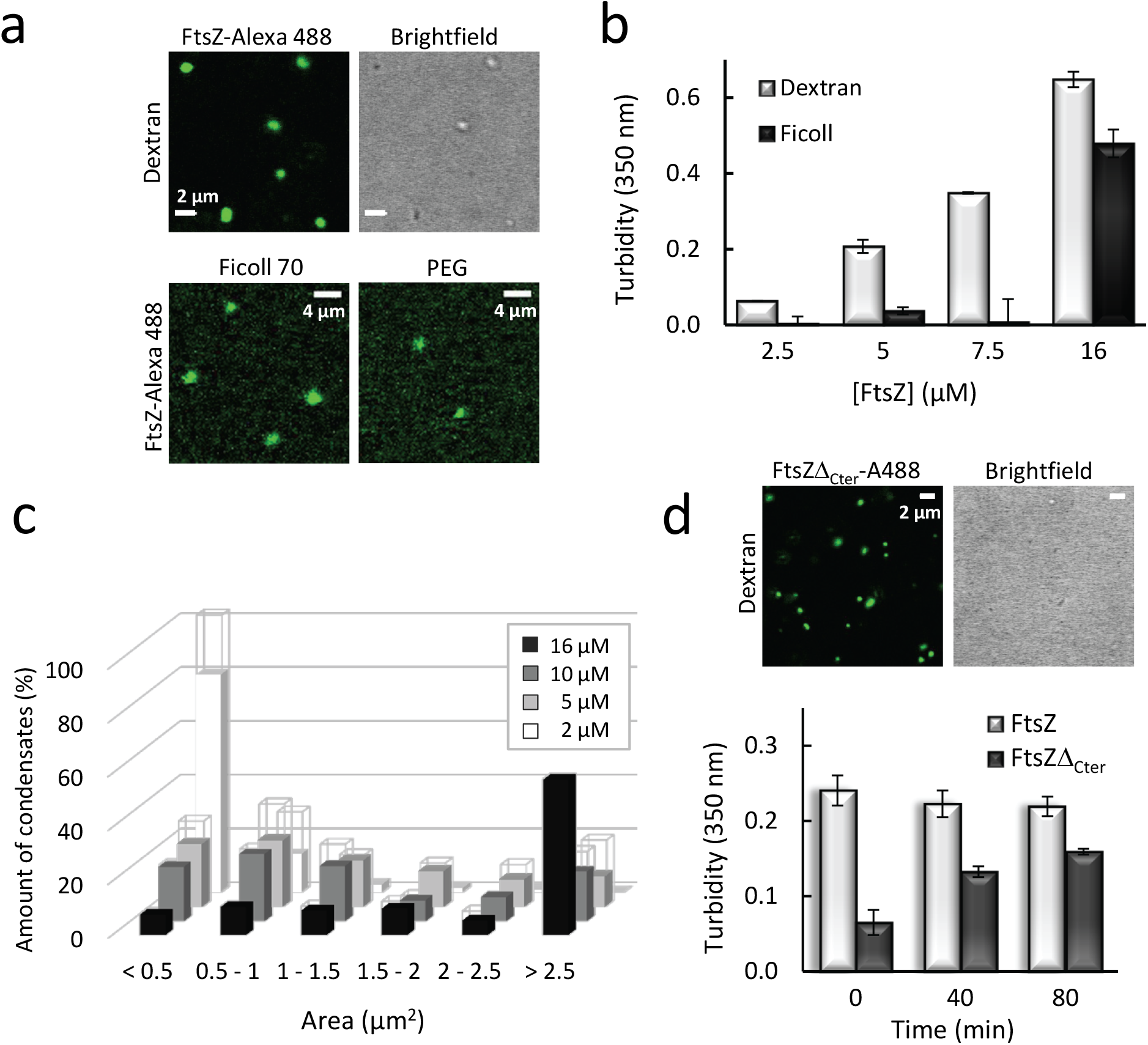
FtsZ forms condensates. (**a**) Confocal microscopy and transmitted images of condensates of FtsZ in different crowders. (**b**) Dependence of turbidity signal on FtsZ concentration in dextran and Ficoll. Note that signal corresponds to all structures present in the solution, including condensates and irregular arrangements. (**c**) Size distribution of condensates in the presence of dextran at various FtsZ concentrations (in ascending order, n = 124, 99, 80 and 106 particles). Errors, depicted as open sections of the bars, correspond to S.D. from 2 (2 and 5 μM FtsZ) or 3 (10 and 16 μM FtsZ) independent images. (**d**) Confocal microscopy and transmitted images of condensates formed by the FtsZ mutant, FtsZΔ_Cter_, in dextran. Below, changes in the turbidity signal of the FtsZ mutant condensates over time, with those of wild-type FtsZ shown as reference. Except when specified, FtsZ and FtsZΔ_Cter_ concentrations were 5 μM. Dextran and Ficoll concentrations were 200 g/l, and PEG was 50 g/l. All experiments in working buffer. In (b) and (d) data are the average of at least 3 independent experiments ± S.D., except for 2.5 and 7.5 μM FtsZ with dextran in (b), measured in duplicate.

Thorough characterization in the presence of dextran showed that the size, shape and abundance of the FtsZ condensates depended on the salt, magnesium and protein concentration. At 100 mM KCl, confocal microscopy and turbidity measurements indicated that reducing Mg^2+^ concentration from 10 mM to 1-5 mM significantly reduced the number of structures, except if protein concentration was tripled to 16 μM (**Supplementary Fig. S2a,b**). At 10 mM Mg^2+^ and high protein concentration, irregular FtsZ assemblies of large size were mostly observed. Decreasing protein concentrations resulted in the progressive appearance of smaller more regular structures at the expense of the irregular arrangements, reaching a population of mostly round structures resembling condensates at and below 5 μM FtsZ (**Fig. 1a** and **Supplementary Fig. S2a**). This was accompanied by a decrease in the size distribution of the condensates within this protein concentration range (**Fig. 1c**), and in the turbidity values arising from all species present in the solution (condensates and/or irregular arrangements) (**Fig. 1b** and **Supplementary Fig. S2b**). An increase in salt to 300 mM KCl resulted in scarce and very small, apparently round, structures, even with high (10 mM) Mg^2+^ content (**Supplementary Fig. S1**), consistent with low turbidity values (**Supplementary Fig. S2c**). Only at high protein concentrations was a significant population of arrangements of variable shape apparent (**Supplementary Fig. S1)**. These results are in line with our previous study on FtsZ∙SlmA∙SBS biomolecular condensates, at 1 mM Mg^2+^, 300 mM KCl, and 150 g/l crowding agents, where FtsZ alone was homogeneously dispersed in solution (Monterroso *et al.*, 2019). Condensation in the presence of Ficoll followed the same trend with KCl and Mg^2+^ concentration as that described with dextran (**Supplementary Fig. S3a,b**).

These experiments show that FtsZ assembles into defined micrometer-sized arrangements in crowding solutions at protein, Mg^2+^ and salt concentrations strongly favoring self-association of the protein monomers. As round structures compatible with biomolecular condensates prevailed at 5 μM FtsZ, in 50 mM Tris-HCl, pH 7.5 with 100 mM KCl, 10 mM MgCl_2_ (working buffer) and 200 g/l dextran, these conditions were selected for their further characterization.

### Condensation of FtsZ lacking the unstructured C-terminal region is less efficient

Unstructured domains of proteins have been implicated as important contributors to biomolecular condensation. To evaluate the impact of such domains on the formation of the condensate-like arrangements described above, we investigated the mutant FtsZΔ315-383 (FtsZΔ_Cter_), which contains the globular domain of FtsZ but lacks the unstructured C-terminal flexible linker as well as the conserved C-terminal peptide known to bind other cell division proteins (Ortiz *et al*, 2016). We found that this mutant FtsZ retained the ability to form condensates in crowded solutions under the experimental conditions described above (**Fig. 1d**). Interestingly, these condensates in the images were substantially smaller than those formed by the wild-type protein at comparable times. This observation is compatible with the lower turbidity values measured for the mutant protein after mixing with the crowder. While the turbidity values remained constant for at least 80 min in the case of the wild-type FtsZ, those for the mutant noticeably increased within this same time interval, suggesting that the condensation process is slower. Analysis of this mutant by analytical ultracentrifugation showed that it retains the ability to form oligomers in solution (**Supplementary Fig. S4**), which may explain the tendency of this protein to form condensates. Together, this evidence suggests that the unstructured domain of FtsZ may not be strictly required to mediate its assembly into condensates, but probably has a role in enhancing the overall process.

### FtsZ condensates are dynamic and evolve into filaments in the presence of GTP

Next, we asked whether the round FtsZ structures resembling molecular condensates were dynamic. To this end, we followed three different approaches: incorporation of added FtsZ on preformed condensates, evolution of condensates over time, and GTP-triggered polymerization of FtsZ forming the condensates.

First, we performed capture experiments using two separate pools of FtsZ labeled with spectrally different dyes, and similarly to that described for other condensates (Monterroso *et al.*, 2019; Woodruff *et al*, 2017). We found that FtsZ-Alexa 488 readily incorporated into preformed condensates containing FtsZ labeled with Alexa 647 (FtsZ-Alexa 647) as a tracer, suggesting that the population of FtsZ within the condensates is dynamic. Images show the initial and final states and the stepwise diffusion of the added protein into the condensates (**Fig. 2a**). Capture experiments using the reverse dyes yielded the same result (**Supplementary Fig. S5a**). Likewise, FtsZΔ_Cter_ condensates proved to be dynamic following the same approach (**Fig. 2b**).

**Figure 2.**
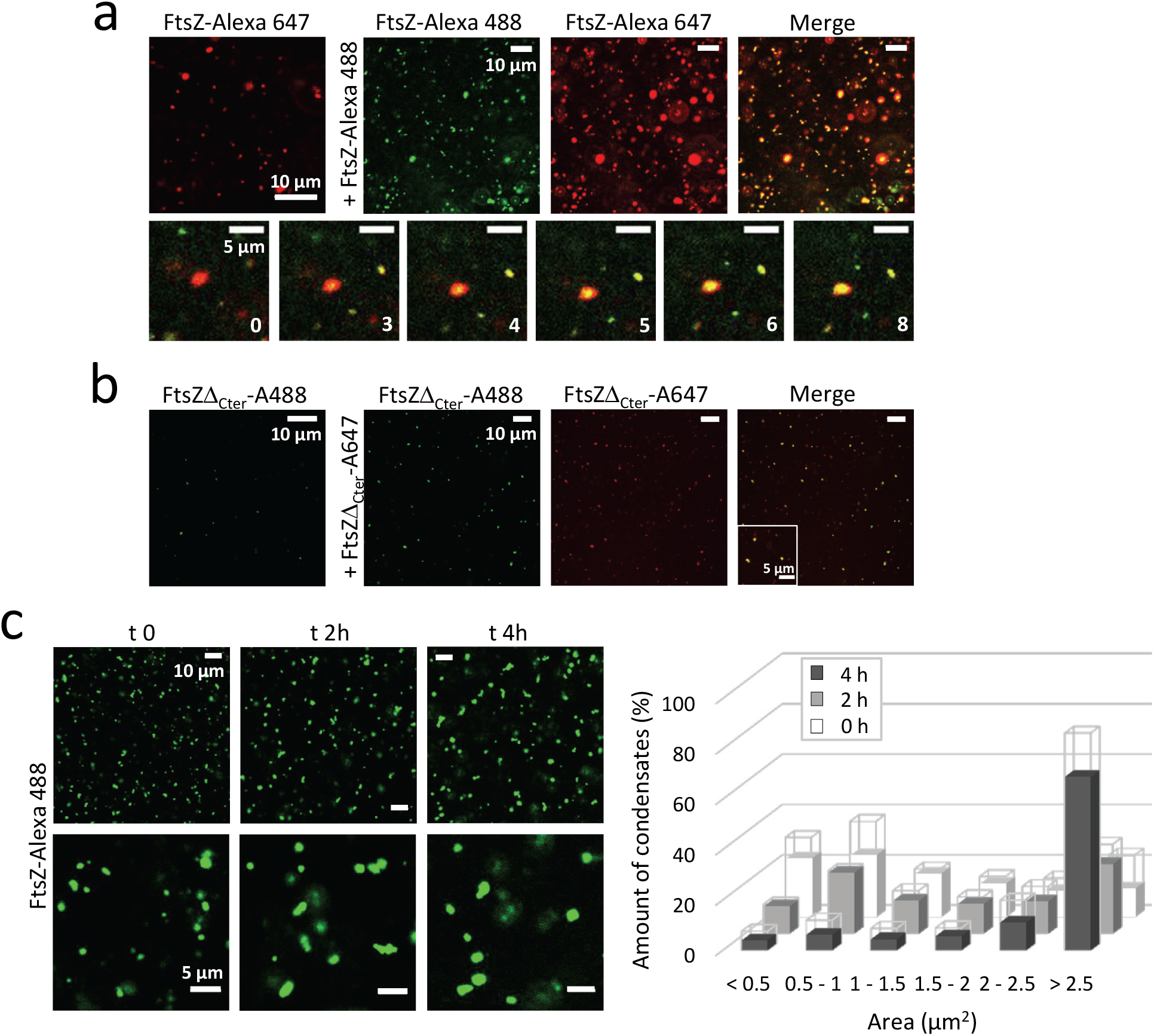
Dynamic nature of FtsZ condensates. (**a**) Representative confocal microscopy images showing the initial and final states after addition of FtsZ-Alexa 488 into FtsZ condensates with FtsZ-Alexa 647 as a tracer and, below, stepwise diffusion at the times indicated in seconds. (**b**) Representative confocal microscopy images showing the initial and final states after addition of FtsZΔ_Cter_-A647 into FtsZΔ_Cter_ condensates with FtsZΔ_Cter_-A488 as a tracer. (**c**) Images of FtsZ condensates with time, and corresponding distribution of sizes on the right (n= 99, 157 and 206 particles for 0, 2 and 4 h, respectively). Errors, depicted as open sections of the bars, correspond to S.D. from 2 (2h) or 3 (0 and 4h) independent images. FtsZ and mutant concentrations were 5 μM. All experiments in working buffer with 200 g/l dextran.

We also monitored the evolution of FtsZ condensates over time. We found a clear shift of the condensate size distribution towards larger values over a 4 h time course (**Fig. 2c**). As no significant background signal (*i.e*. no free protein) was observed in the images, this is indirect evidence of fusion among different condensates, which has been previously observed in other condensate assemblies.

Lastly, we tested the responsiveness of the FtsZ condensates to GTP, which is well known to induce FtsZ polymer formation and is a hallmark of its functionality *in vivo*. Confocal images showed that addition of GTP strongly induced the polymerization of FtsZ that seemed to occur with a concomitant reduction in the number and/or size of the condensates, although both structures coexisted during the time window monitored (**Fig. 3a**). Further insight into the behavior of the condensates upon GTP supply was obtained by turbidity measurements, showing a significant decrease in the signal arising from the condensates immediately after nucleotide addition (**Fig. 3b**). Moreover, measurements with condensates formed using Ficoll as crowding agent exhibited the same behavior (**Supplementary Fig. S3c**). This, together with the incipient polymer formation observed in the images taken at short times, strongly suggests that the polymers originate from the condensates. Had polymerization exclusively come from any possible unassembled protein coexisting with irreversible condensates, unlikely given the negligible background signal observed in the images under these conditions, we would have detected an increase in turbidity instead of the strong decrease observed. Monitoring these samples over longer times showed that the turbidity signal continued to slightly decrease up to a point, dependent on the nucleotide concentration, above which it gradually recovered, reaching a value close to that in the absence of GTP (**Fig. 3b** and **Supplementary Fig. S6a**). This behavior is consistent with the characteristic dissociation of the FtsZ polymers due to GTP exhaustion (Mukherjee & Lutkenhaus, 1998).

**Figure 3.**
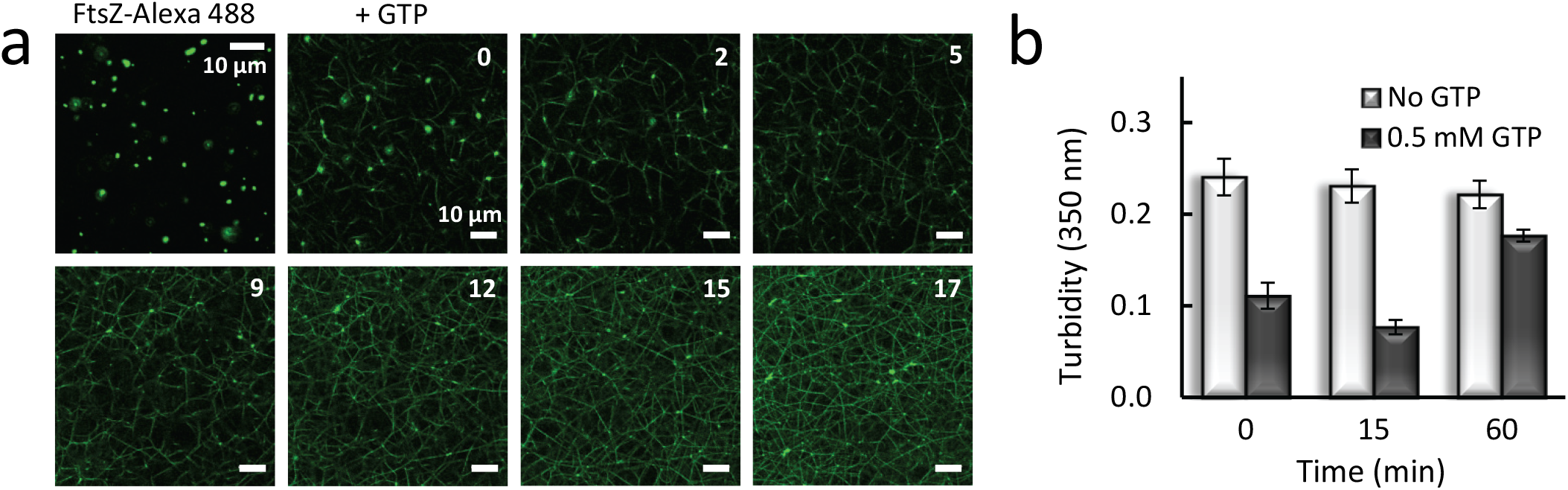
Functionality of FtsZ within the condensates. (**a**) Representative confocal images of FtsZ condensates before polymerization and evolution after GTP (0.5 mM) addition at the indicated times, in minutes. All images of the time lapse correspond to a single field. (**b**) Evolution of the turbidity signal of condensates with time and variation after GTP addition at time zero. Values correspond to the average of 3 independent experiments ± S.D. All experiments were conducted with 5 μM FtsZ in working buffer with 200 g/l dextran.

Interestingly, when conducting the experiments in conditions under which the round condensates coexisted with more irregular forms, the latter also exhibited dynamic protein capture and conversion into the typical GTP-triggered FtsZ polymers, as monitored by confocal microscopy (**Supplementary Figs. S5b** and **S6b,c**). Addition of GTP to these samples rendered a decrease in the turbidity signal, as observed with solutions where the round condensates are the major species (**Supplementary Fig. S6d**).

### Reconstruction of FtsZ condensates in cytomimetic microdroplets

To determine whether FtsZ condensates could also be formed in a confined cell-like system, we used microfluidics to encapsulate the protein (**Fig. 4**). Confocal microscopy images of FtsZ, containing FtsZ-Alexa 488 as a tracer, encapsulated in microdroplets stabilized by an *E. coli* lipid mixture boundary, indicated the presence of condensates in the z-sections of the microdroplets with no apparent preference for the lumen or the lipid interface. The presence of these structures was more obvious in the maximal projection of the images (far right in **Fig. 4a**). No background signal was observed in the lumen, while a strong fluorescence demarcating the lipid boundary was found.

**Figure 4.**
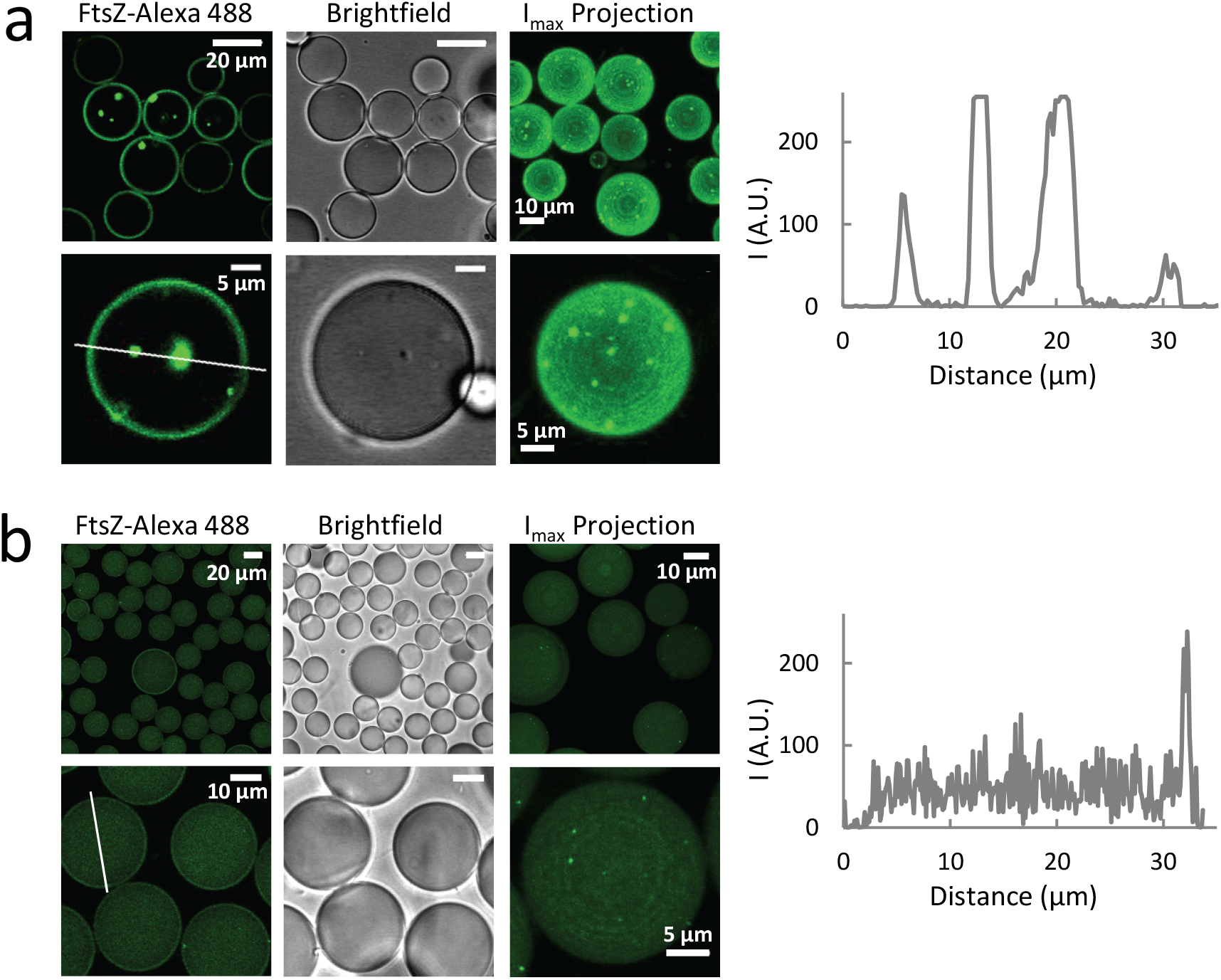
Condensates of FtsZ encapsulated in microdroplets stabilized by the *E. coli* lipid mixture. (**a**) Confocal microscopy images of microdroplets generated by microfluidics containing the condensates. Experiments were with 5 μM FtsZ in working buffer with 200 g/l dextran. (**b**) Encapsulation under experimental conditions not promoting condensation of FtsZ in bulk: 12 μM FtsZ in 50 mM Tris-HCl, pH 7.5, 1 mM MgCl_2_, 300 mM KCl and 150 g/l dextran. The third column of images for (a) and (b) are maximum intensity projections corresponding to different fields. To the right of (a) and (b) are intensity profiles of the green channel obtained across the lines drawn in the respective images.

Encapsulation of FtsZ under conditions that discourage bulk formation of condensates or any other detectable structures (300 mM KCl, 1 mM MgCl_2_, 150 g/l dextran) showed scarce and very small condensates in the z-sections, which were more obvious in the maximal projection and mostly near the membrane (**Fig. 4b**). Under these conditions, the protein remained homogeneously distributed within the lumen, in good agreement with previous reports showing FtsZ encapsulated under similar conditions (Mellouli *et al*, 2013; Sobrinos-Sanguino *et al*, 2017).

These experiments confirm that condensates are still assembled when FtsZ is encapsulated under conditions favoring their formation in bulk. Moreover, incipient condensate formation was also apparent when the protein was encapsulated in conditions under which no condensates were detected by confocal microscopy or turbidity, strongly suggesting that confinement and/or the lipid membrane, directly or indirectly, influence their formation.

## Discussion

Here we show that purified FtsZ protein forms structures compatible with biomolecular condensates, observed both in bulk solutions and when reconstructed in cytomimetic platforms. These structures are mostly reservoirs of material that have exceeded their solubility in the phase outside the condensate. Their assembly, disassembly, abundance and coexistence with irregular, but still active, clusters are influenced by crowding and other chemical conditions that affect oligomerization of GDP-bound FtsZ (**Fig. 5a**). This behavior is probably derived from the well-known self-association properties of this protein, which provide the multivalent states that typically favor formation of condensates.

**Figure 5.**
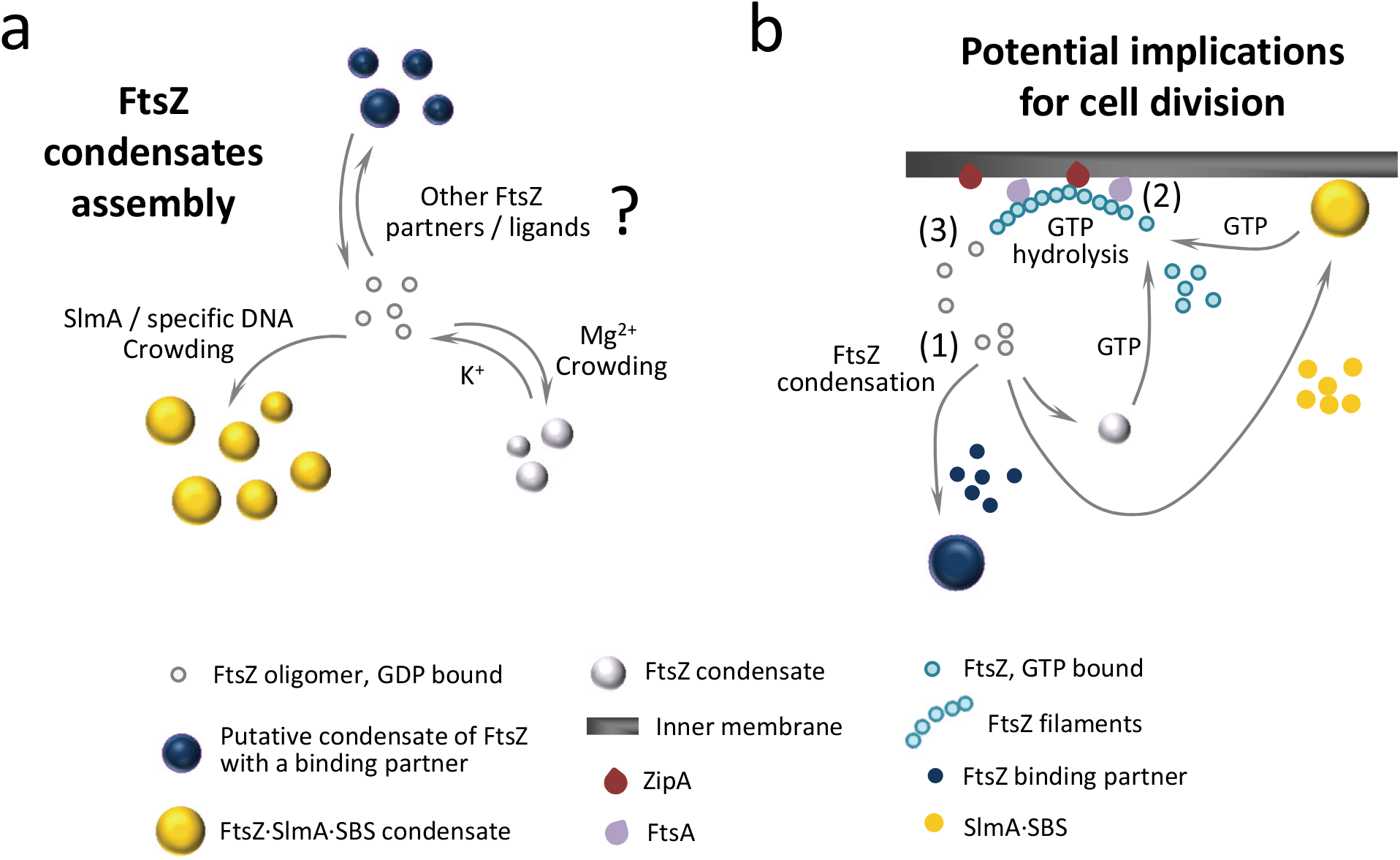
Formation of FtsZ biomolecular condensates and potential role in bacterial cell division regulation. (**a**) FtsZ condensate assembly is disfavored by K^+^, favored by crowding and Mg^2+^ and strongly promoted by the SlmA·SBS nucleoprotein complex. It is possible that condensation is also positively or negatively regulated by additional binding partners or by natural or synthetic ligands of FtsZ. (**b**) In the cell, FtsZ oligomers could form condensates on their own or together with the nucleoid occlusion factor SlmA (FtsZ·SlmA complexes tend to locate at the cellular membrane (Monterroso *et al.*, 2019)), and possibly also with other division regulators (1). In the presence of GTP, FtsZ would leave the condensates and associate into filaments attached to the bacterial membrane through the natural anchors, ZipA and FtsA (2). GTP hydrolysis would lead to GDP-bound FtsZ subunits within the filaments, and a loss of their longitudinal interactions would result in the release of FtsZ (3). This released FtsZ would be able to reassemble into homotypic or heterotypic dynamic condensates, contributing to the spatiotemporal regulation of bacterial cell division.

FtsZ within the condensates remained responsive to ligands, only forming filaments upon addition of GTP. This GTP-dependent shuttling of FtsZ between condensate and polymer states was also observed in our previous study of FtsZ·SlmA complexes, which showed that when mixed with the DNA-binding SlmA protein under crowding conditions, FtsZ formed condensates capable of reversible evolution to polymers in the presence of GTP (Monterroso *et al.*, 2019). The present work and our previous study on FtsZ·SlmA condensates emphasize the key role of GDP in the formation of FtsZ condensates. It should be noted that FtsZ behaves differently from other NTP-bound proteins such as DEAD-box ATPases that partition to the condensed phase, with ATP hydrolysis promoting disassembly of the condensates (Hondele *et al*, 2019).

FtsZ condensate formation is reversible, regulated by ion concentrations and salt, as expected for assemblies involving electrostatic forces (without excluding other possible types of interactions), and that achieve sufficient multivalency to form condensates under conditions that enhance FtsZ oligomerization (*i.e*. low ionic strength and high Mg^2+^ (**Fig. 5a**)). A remarkable difference between FtsZ condensates and those previously described for FtsZ·SlmA·SBS (Monterroso *et al.*, 2019) is that the latter assemble under a much wider variety of conditions. This is consistent with the higher multivalency conferred on the system by the presence of SlmA, a dimer that binds in pairs to a specific SBS site on DNA (Cabre *et al*, 2015), while also establishing contacts with the C-terminal tail and the folded domain of FtsZ (Du & Lutkenhaus, 2014).

Protein condensate formation is usually associated with regions of low sequence complexity (Banani *et al.*, 2017) and, indeed, FtsZ contains a ~50 residue flexible unstructured segment (Gardner *et al*, 2013). The mutant FtsZ comprising the globular region of the protein but not the linker shows condensation, possibly related to the multivalency emerging from its oligomerization interfaces. Nonetheless, this mutant displays slower condensation than wild-type FtsZ, which might be a result of differences in their dynamics and/or topology. Along these lines, GTP-induced filaments of FtsZ mutants from different bacterial species lacking the C-terminal peptide and/or the linker exhibited morphological alterations and slower polymerization rates (Buske & Levin, 2013). Also, as here with FtsZ, examples can be found where condensation is observed for folded domains, with disordered regions playing a modulatory role (Kroschwald *et al*, 2018; Riback *et al*, 2017).

We observed formation of FtsZ condensates in confined cell like environments that included crowding and a membrane boundary. The use of such synthetic systems with controlled composition simplifies the interpretation of condensation compared with *in vivo* studies and, in the particular case of bacteria, circumvents technical obstacles due to their small size (Azaldegui *et al.*, 2020). Our encapsulation studies indicate that fully formed FtsZ condensates do not show a preference for the membrane, although it seems that the membrane could participate in the condensation of FtsZ, as shown by the small emerging structures at the lipid surface under bulk non-condensation conditions. This is compatible with either an increase in size of submicrometer condensates already present in solution, providing a surface for nucleation (Snead & Gladfelter, 2019), and/or further enhancement of their formation by confinement within the microdroplet. Interestingly, FtsZ·SlmA nucleoprotein condensates do accumulate at the lipid boundary (Monterroso *et al.*, 2019), probably because of SlmA, which we recently found to interact with membranes (Robles-Ramos *et al*, 2020).

How might biomolecular condensation of FtsZ be relevant *in vivo*? Under certain conditions, cellular FtsZ oligomers might assemble into condensates, either alone or combined with any of FtsZ’s multiple binding partners (**Fig. 5b**). Condensate formation would be modulated to some extent by weak, transient electrostatic interactions with the environment, as nucleic acids are abundant and most proteins in *E. coli* are polyanions at intracellular pH (Spitzer & Poolman, 2009). As condensation enables dynamic spatial localization of cellular processes, condensates containing FtsZ might be favored during a particular phase of the cell cycle, perhaps used as a cytoplasmic storage form, or as a response to stresses (see below). For instance, the local accumulation of a higher number of molecules provided by condensation could permit more rapid pre-assembly with key protein partners or ligands than by depending on successive recruitment of cell division proteins in the cytoplasm or on the cytoplasmic membrane. In the presence of GTP, FtsZ would exit the condensates and associate into mobile complexes of filaments attached to the bacterial membrane through the natural anchor proteins, ZipA and FtsA (Baranova *et al*, 2020; Rowlett & Margolin, 2014; Yang *et al*, 2017). GTP hydrolysis would increase the amount of FtsZ subunits bound to GDP within the filaments, with the associated loss in their longitudinal interactions and subsequent release of FtsZ (Du & Lutkenhaus, 2019), now available for a new condensation cycle. Curiously enough, the elusive ultrastructure of the *E. coli* Z-ring, only recently revealed through high resolution imaging, shows loosely associated clusters of FtsZ molecules of round appearance (Du & Lutkenhaus, 2019; Lyu *et al*, 2016). Further work will be required to ascertain the interplay of the FtsZ condensates identified here or of other putative heterotypic condensates involving this protein with the division ring.

The biomolecular condensation of purified FtsZ demonstrated here adds to the growing number of examples of this type of behavior by bacterial proteins. Despite initial doubts raised by the lower frequency of intrinsically disordered regions in bacterial proteomes, very recent research suggests that biomolecular condensation is an organizational principle that also operates in these microorganisms (Azaldegui *et al.*, 2020). As an alternative to the membranous organelles of eukaryotes, such condensates distributed within the bacterial cytoplasm might contribute to the spatial regulation of various metabolic reactions, a role once attributed almost exclusively to the bacterial membrane envelope. In particular, condensates might form preferentially under stress conditions. Interestingly, it was recently reported that late stationary phase *E. coli* assemble protein aggregates at their cell poles containing FtsZ and other proteins; these “regrowth delay bodies” dissolve once growth is resumed, suggesting that they are dynamic (Yu *et al*, 2019). Such bodies may consist of multiple proteins that tend to form condensates under certain stress conditions such as starvation, desiccation and persister states (Laskowska & Kuczynska-Wisnik, 2020). It is noteworthy that cells in such states are likely deficient in GTP, a situation that as suggested by our results, could contribute to the formation of FtsZ condensates. Given that FtsZ is considered a promising target in the quest for new antibiotics (Kusuma *et al*, 2019), and that cells in the persister state are particularly resistant to antibiotics (Fisher *et al*, 2017), understanding how FtsZ forms condensates and where and when such condensates might form in cells may provide additional clues to fight against antimicrobial resistance.

## Methods

### Reagents

GTP, dextran 500 (500 kDa), PEG 8 (8 kDa) and other analytical grade chemicals were from Sigma Chemical Co., St. Louis MO, USA. Ficoll 70 (70 kDa) was from GE Healthcare, IL, USA. Crowders were dialyzed in 50 mM Tris-HCl, 100 or 300 mM KCl, pH 7.5, and their concentration measured as earlier described (Monterroso *et al*, 2016). Polar extract phospholipids from *E. coli* (Avanti Polar Lipids, Alabaster AL, USA), were stored in chloroform at −20 °C. Shortly before use they were thoroughly dried in a Speed-Vac device and the resulting film resuspended in mineral oil by two cycles of vortex and 15 min sonication in a bath. Final concentration of the lipids in mineral oil was 20 g/l.

### FtsZ and FtsZΔ_Cter_ expression, purification and labeling

Wild-type *E. coli* FtsZ was isolated as described elsewhere (Rivas *et al.*, 2000) and stored at −80 °C until used. The plasmid expressing the FtsZΔ_Cter_ mutant (FtsZΔ315-383, which lacks residues comprising the unstructured region and the C-terminal tail of FtsZ) was kindly provided by P. Schwille (Max Planck Institute of Biochemistry, Martinsried) and purified following the same procedure. Covalent labeling of amine groups of the proteins with Alexa Fluor 488 or Alexa Fluor 647 succinimidyl ester dyes (Molecular probes/Invitrogen) was conducted in its polymer-assembled form as previously described (González *et al*, 2003; Reija *et al*, 2011). The labeling ratio, calculated from the molar absorption coefficients of the proteins and the dyes, ranged between 0.1-0.9 moles of dye per mole of protein.

### Preparation of bulk samples and selection of final conditions

FtsZ was directly added to solutions containing the crowders at the specified conditions of KCl and magnesium in 50 mM Tris-HCl, pH 7.5, and incubated for ~ 30 min. When required, polymerization was triggered by diffusion of GTP directly added over the mixture. For imaging experiments, labeled proteins were used as tracers (final concentration 0.5 or 1 μM, < 10% of total protein concentration). Images were acquired with different dyes, both for wild-type (FtsZ-Alexa 488, FtsZ-Alexa 647) and mutant proteins (FtsZΔ_Cter_-A488, FtsZΔ_Cter_-A647) with equivalent results. Unless otherwise specified, experiments were conducted with 5 μM FtsZ in working buffer (50 mM Tris-HCl pH 7.5, with 100 mM KCl, 10 mM MgCl_2_) and 200 g/l dextran.

### Microfluidic chip fabrication and FtsZ encapsulation

The devices used were constructed by conventional soft lithographic techniques from masters kindly provided by the W.T.S. Huck group (Radboud University, Nijmegen, The Netherlands; chip design detailed elsewhere (Mellouli *et al.*, 2013)) following the procedure described (Monterroso *et al.*, 2019).

Encapsulation was conducted by mixing, in a 1:1 ratio prior to the droplet formation junction, two streams of dispersed aqueous phases containing FtsZ-Alexa 488 (0.5 μM) as a tracer and, except when stated, FtsZ (5 μM) in working buffer with 200 g/l dextran. The third stream supplied the *E. coli* lipid mixture at 20 g/l in mineral oil. Data presented correspond to experiments delivering solutions at 150 μl/h (oil phase) and 20 μl/h (total aqueous phases) by automated syringe pumps (Cetoni GmbH, Germany) yielding uniform droplets with average diameters of 22 μm. Production of the lipid microdroplets in the microfluidic chip was monitored using an Axiovert 135 fluorescence microscope (Zeiss).

### Diffusion of additional FtsZ into preformed condensates

Samples with FtsZ at the specified final concentration containing 0.5 μM FtsZ labeled with Alexa 647 as a tracer were prepared and imaged before and after addition of 0.5 μM FtsZ-Alexa 488. Experiments adding FtsZ-Alexa 647 to condensates labeled with FtsZ-Alexa 488 as a tracer were also conducted. The diffusion of the added labeled protein into the condensates was monitored over time. Experiments with the mutant were done following the same procedure.

### Confocal microscopy measurements and data analysis

Condensates and microfluidics microdroplets were visualized in silicone chambers (Molecular probes/Invitrogen) glued to coverslips. Images were obtained with Leica TCS-SP2 or TCS-SP5 inverted confocal microscopes with a HCX PL APO 63× oil immersion objective (N.A. = 1.4; Leica, Mannheim, Germany). 488 and 633 nm laser excitation lines were used to excite Alexa 488 and Alexa 647 dyes, respectively. Several images were registered across each sample, corresponding to different observation fields. Brightfield and fluorescence images were taken simultaneously. ImageJ (National Institutes of Health, USA) was used to produce images and, after noise reduction by applying a Kuwahara filter and threshold correction by visual inspection, to measure the areas distribution of FtsZ condensates with the particle analysis option of the software.

### Turbidity measurements

Turbidity of samples (125 μl solutions) aimed at determining the effect of crowders, salts or FtsZ concentration on condensate formation, and their response to GTP addition, was recorded at room temperature and 350 nm using a Varioskan Flash plate reader (Thermo Fisher Scientific, MA, USA). The absorbance was measured after 30 min incubation. For time dependence measurements, data were taken every 5 min for 90 min. Reported values are the average of at least 3 independent experiments ± S.D., unless otherwise stated.

## Data availability

The datasets generated and/or analyzed during the current study are available from the corresponding authors upon reasonable request.

## Acknowledgements

We thank H. Yébenes for advice on, and preliminary, image analysis and M. Sobrinos-Sanguino for technical support. We also thank the staff of CIB Margarita Salas Confocal Laser and Multidimensional Microscopy (M.T. Seisdedos and G. Elvira) and Molecular Interactions (J.R. Luque) Facilities for excellent assistance in imaging and ultracentrifugation experiments, and the Technical Support Facility for invaluable input. This work was supported by the Spanish Ministerio de Economía y Competitividad (BFU2014-52070-C2-2-P and BFU2016-75471-C2-1-P, AEI/FEDER, UE, to G.R.), by the Spanish Ministerio de Ciencia e Innovación (2019AEP088 and PID2019-104544GB-100, AEI, to G.R. and S.Z.), and by the National Institutes of Health (GM131705, to W.M.). M.Á.R.-R. was supported by the Agencia Estatal de Investigación and the European Social Fund through grant BES-2017-082003. The funders had no role in study design, data collection and interpretation, or the decision to submit the work for publication.

## Author contributions

**MARR**: lead investigation, data curation, supporting formal analysis. **SZ**: conceptualization, supervision, supporting project administration, data curation, validation, investigation, supporting funding acquisition, writing - review & editing. **CA**: supporting investigation, supporting formal analysis. **WM**: conceptualization, writing - review & editing, supporting funding acquisition. **GR**: writing - review & editing, conceptualization, supervision, funding acquisition. **BM**: project administration, conceptualization, supervision, microfluidics methodology, data curation, formal analysis, validation, investigation, writing - original draft, review & editing. **All authors** discussed the results, read and approved the final version of the manuscript.

## Additional Information

### Competing financial interests

The authors declare no competing financial interests.

## Supporting Information

**Supplementary Fig. S1.**
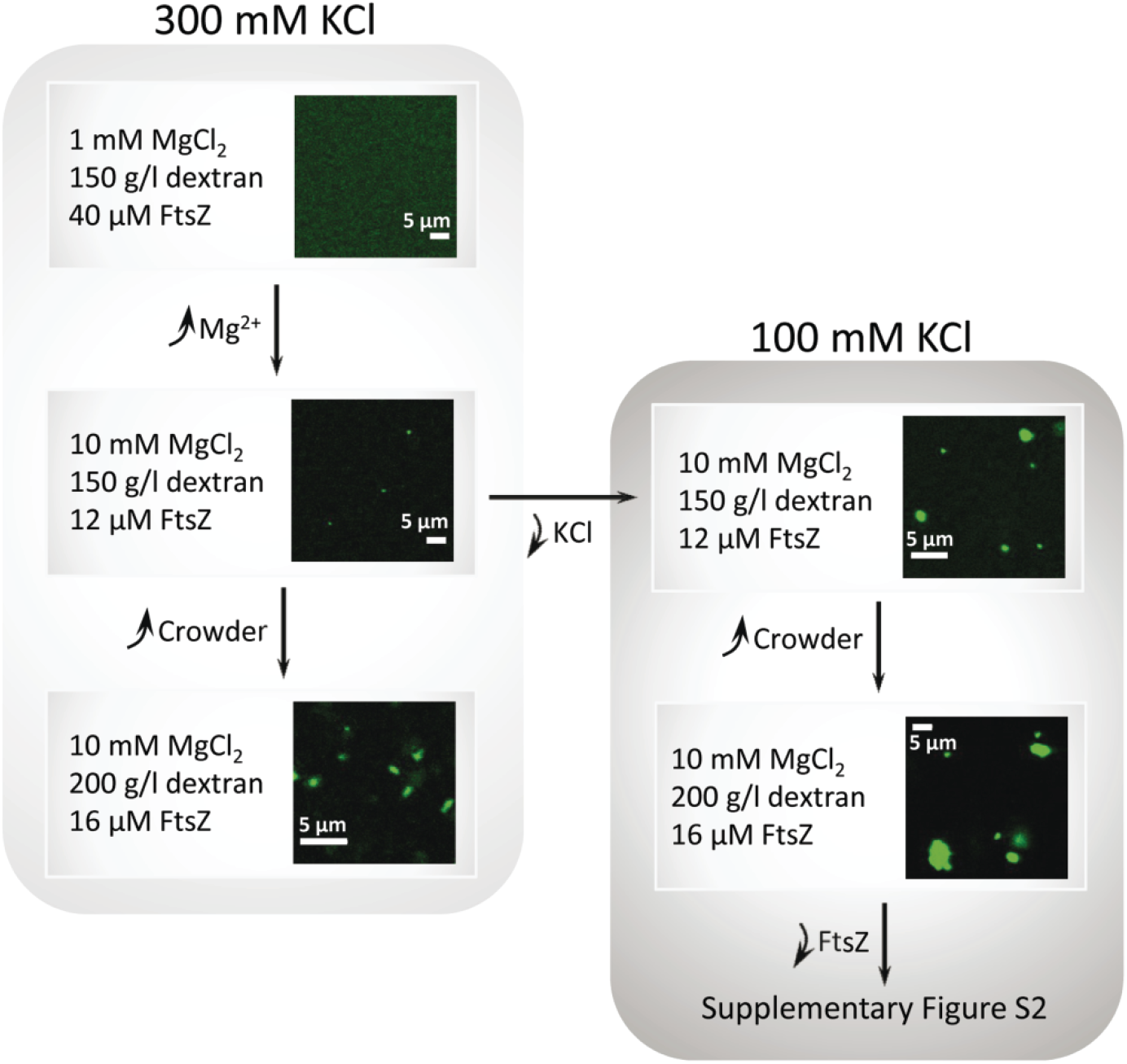
Flow chart of the experimental steps towards the identification of FtsZ condensates, together with representative confocal images of each condition. FtsZ labeled with Alexa 488 as a tracer. Top panel at 300 mM KCl shows the starting condition with no condensates, from our previous study (Monterroso *et al*, 2019).

**Supplementary Fig. S2.**
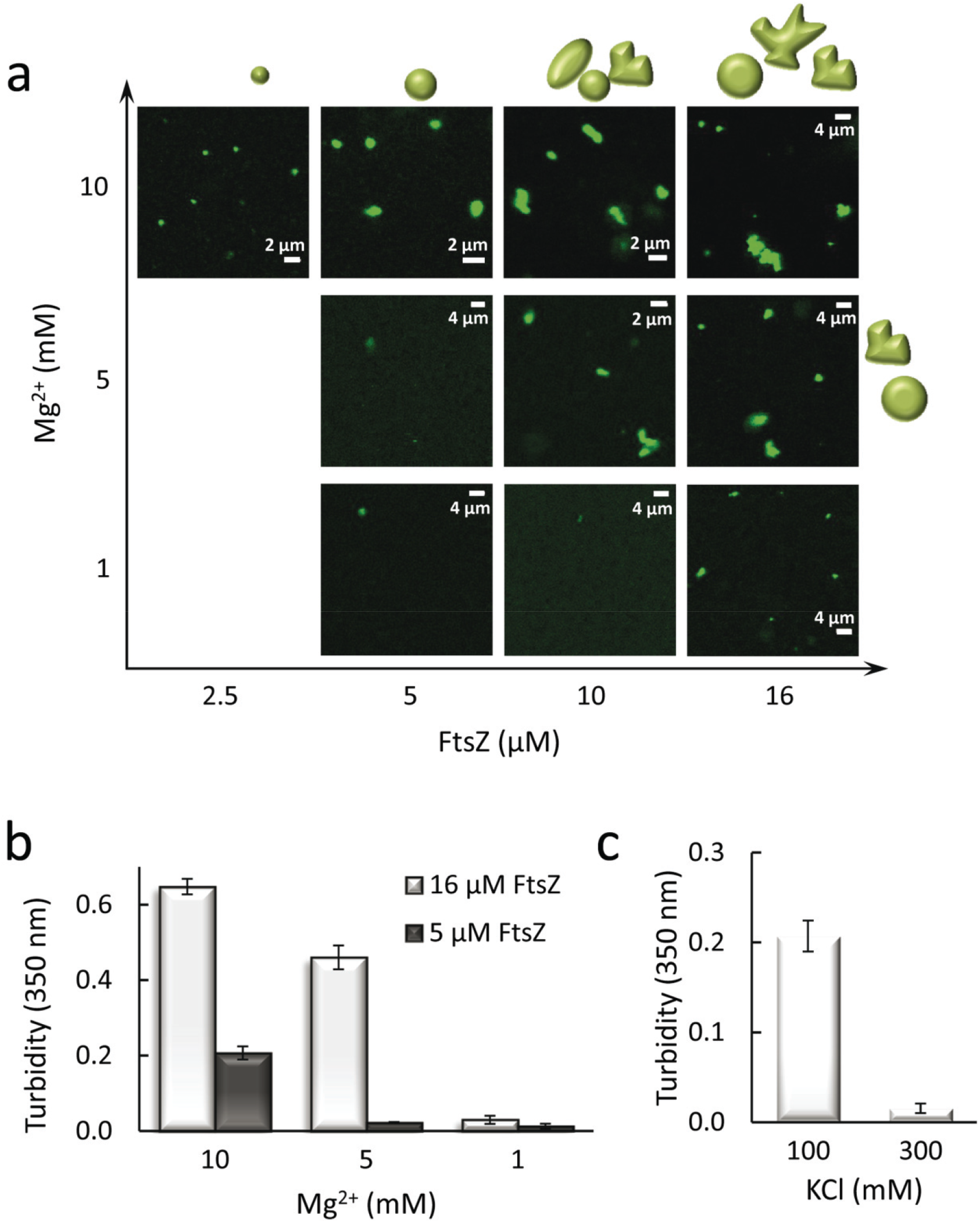
Dependence of the formation of FtsZ arrangements with magnesium, protein and KCl concentration in 200 g/l dextran. (**a**) Representative confocal images of the structures formed by FtsZ, with FtsZ-Alexa 488 as a tracer, at the specified magnesium and protein concentrations. (**b**) Variation in the turbidity signal with magnesium at two FtsZ concentrations. In (a) and (b), KCl was 100 mM. (**c**) Variation in the turbidity signal of FtsZ (5 μM) with KCl concentration at 10 mM magnesium.

**Supplementary Fig. S3.**
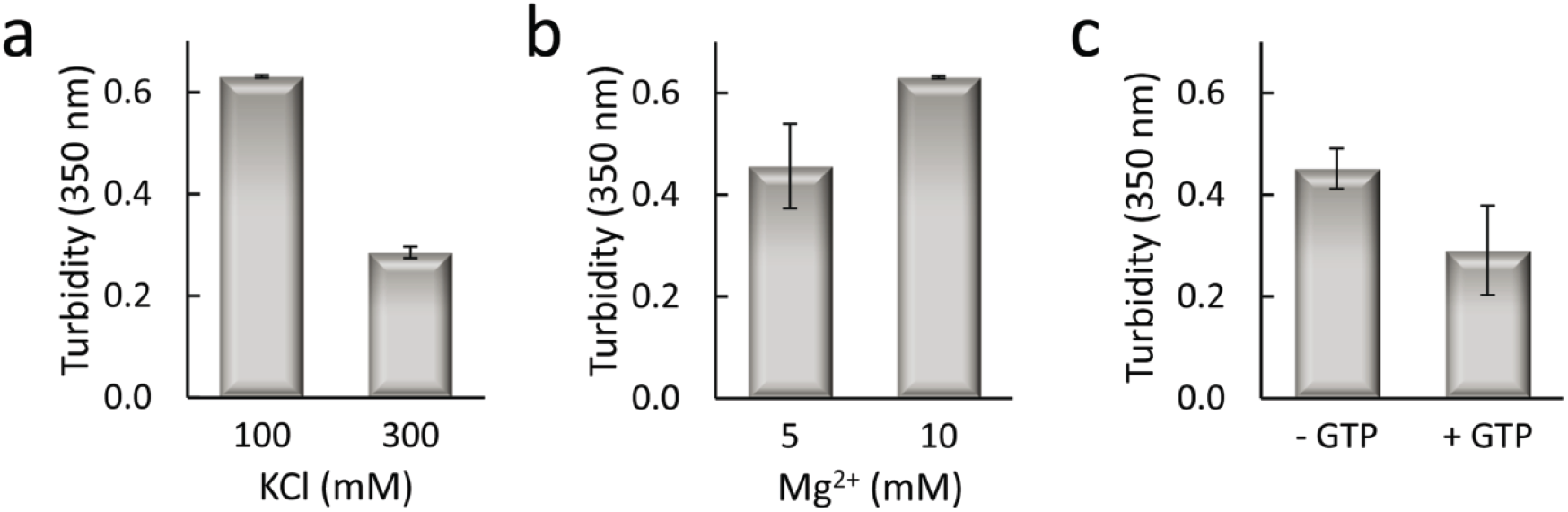
Formation of FtsZ condensates in Ficoll under different experimental conditions. Variation in the turbidity signal of FtsZ in 250 g/l Ficoll with KCl (**a**, 10 mM MgCl_2_) and magnesium concentrations (**b**, 100 mM KCl). (**c**) Effect of 0.5 mM GTP addition on the turbidity signal of FtsZ in working buffer with 200 g/l Ficoll. In all experiments, FtsZ concentration was 16 μM. Note that signals correspond to all structures present in the solution, condensates and/or irregular arrangements depending on the case.

**Supplementary Fig. S4.**
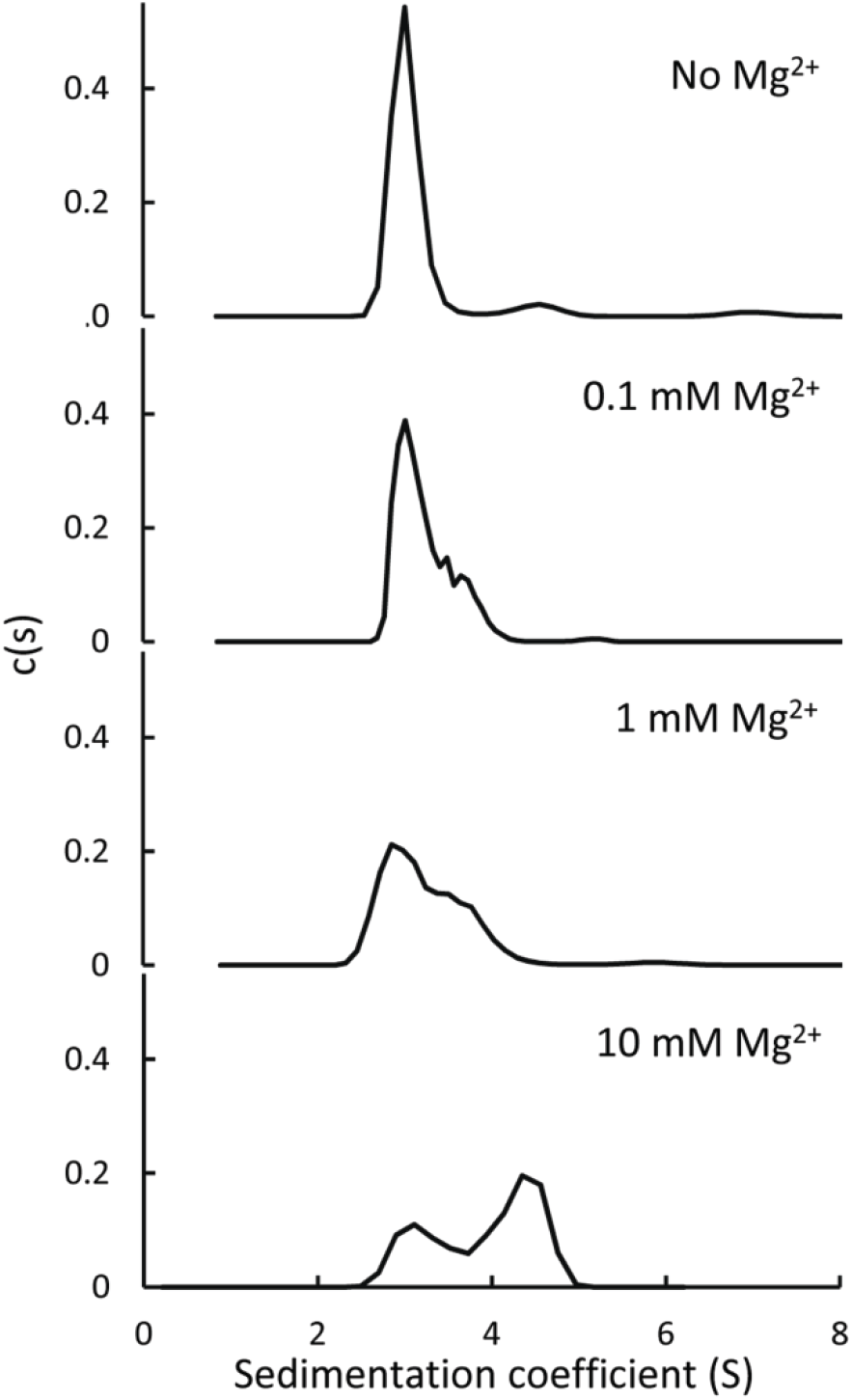
Effect of magnesium on the self-association of FtsZΔ_Cter_ as determined by analytical ultracentrifugation. Shown are sedimentation coefficient distributions of the mutant (15 μM) in 50 mM Tris-HCl pH 7.5, 300 mM KCl, with the specified magnesium concentrations. **Sedimentation velocity experiments of FtsZ mutant.** Experiments were conducted in a XL-I analytical ultracentrifuge (Beckman-Coulter Inc.) equipped with both UV-VIS and Raleigh interference detection systems, using an An-50Ti rotor and 12 mm double sector centerpieces. Sedimentation profiles of samples containing FtsZΔ_Cter_ centrifuged at 48000 rpm and 20 °C were recorded at 260 nm. Concentration of the mutant and experimental conditions are specified in the legend of the figure. Sedimentation coefficient distributions were calculated by least squares boundary modelling of sedimentation velocity data using the c(s) method with SEDFIT (Schuck, 2000). Experimental sedimentation coefficient values (s) were corrected to standard conditions using the program SEDNTERP (Laue, 1992) to obtain the corresponding standard s values (s_20,w_).

**Supplementary Fig. S5.**
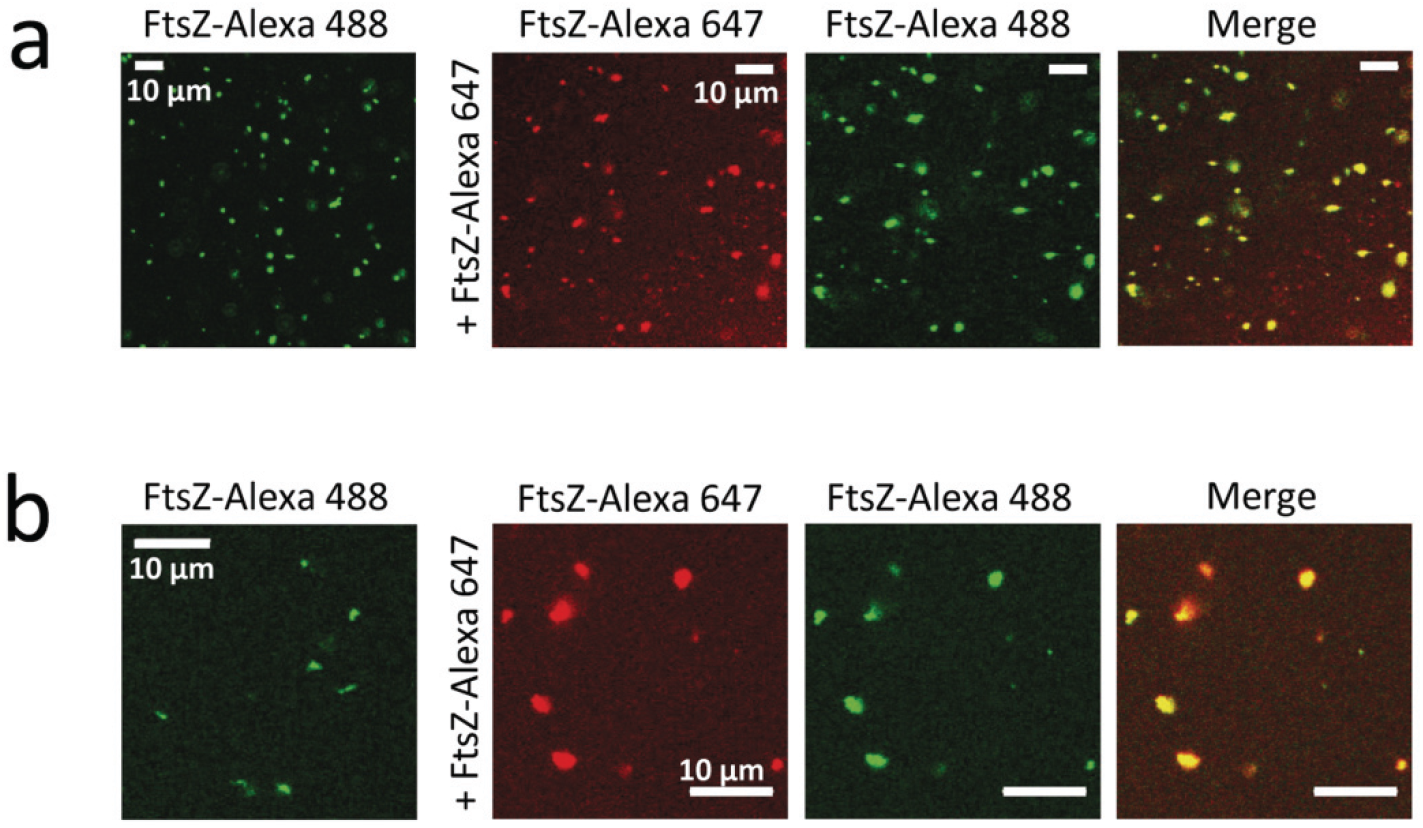
Representative confocal images showing the dynamism of FtsZ condensates under different experimental conditions. Initial and final states after addition of FtsZ-Alexa 647 into FtsZ condensates with FtsZ-Alexa 488 as a tracer in (**a**) 200 g/l dextran and 10 mM MgCl_2_ (5 μM FtsZ) and (**b**) 150 g/l dextran and 5 mM MgCl_2_ (16 μM FtsZ). Both experiments in 50 mM Tris-HCl pH 7.5, 100 mM KCl.

**Supplementary Fig. S6.**
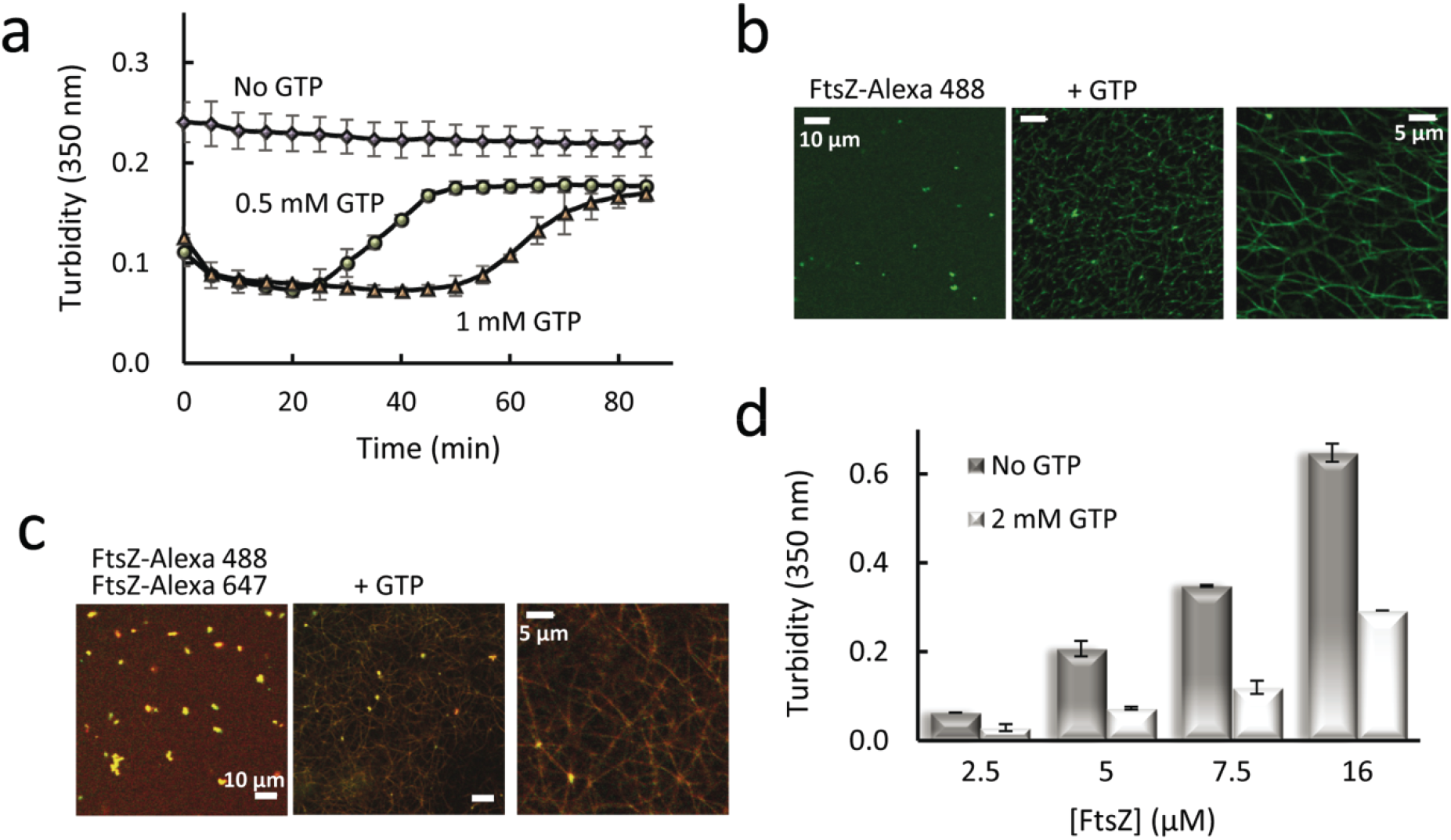
FtsZ within the condensates under different experimental conditions remains active for polymerization. (**a**) Variation in the turbidity signal of FtsZ upon triggering polymerization by addition of the specified GTP concentrations. Evolution of the signal in the absence of GTP is shown for reference. (**b**) Representative confocal images of FtsZ condensates before polymerization and after addition of GTP (1 mM). Image in the far right shows a higher magnification. (**c**) FtsZ condensates after the capture of FtsZ labeled with a complementary dye before polymerization and after addition of GTP (0.75 mM). Image in the far right shows a higher magnification. (**d**) Effect of GTP addition on the turbidity signal of FtsZ at different concentrations of the protein. Note that signal corresponds to all structures present in the solution, condensates and also irregular arrangements depending on the case. Concentrations were 200 g/l dextran, 10 mM MgCl_2_ and 5 μM or the specified FtsZ concentrations (a and d), or 150 g/l dextran, 5 mM MgCl_2_ and 16 μM FtsZ (b and c). All experiments in buffer with 100 mM KCl.

